# Cross-species analysis of experimental PH at single cell resolution reveals prominent contributions *Ednrb*^+^ EC and *Dhcr24*^+^ macrophage populations

**DOI:** 10.1101/2025.04.30.651587

**Authors:** Bolun Li, Yanjiang Xing, Yitian Zhou, Xingqi Xiao, Ting Shu, Xiaomin Song, Hongmei Zhao, Xiaojian Wang, Junling Pang, Peiran Yang, Paul B. Yu, Jing Wang, Chen Wang

## Abstract

**Background:** Animal models are used widely to study pulmonary hypertension (PH). The cell populations that respond to disease-inducing stimuli in these models and their relationship to human disease remain incompletely defined.

**Materials and methods:** This study analyzed the relationship between several rodent models of PH and human disease at single-cell resolution. scRNA-seq was performed on lungs from mice exposed to hypoxia or Sugen/hypoxia, rats exposed to monocrotaline, and controls. A cross-species single-cell dataset was integrated with human lung cell atlas (HLCA) and single-cell dataset from idiopathic pulmonary arterial hypertension (IPAH) to identify overlapping cell subsets between experimental and human disease and species.

**Results:** High levels of overlap were found between species and models of PH, HLCA, and IPAH datasets. Cell subsets perturbed in rat and mouse PH were similar to those found in human disease, with macrophages and endothelial cells being most affected. A novel *Dhcr24^high^* macrophage subset harboring both tissue-remodeling and pro-inflammatory features was consistently increased across models. Several functionally diverse endothelial subtypes were found, including novel Ednrb^+^ and Nox2^+^ subpopulations, reflecting enhanced apoptosis, dysregulated angiogenesis and proliferation, and reactive oxygen species-mediated stress. These macrophage and endothelial subtypes expressed numerous PH drug target genes, and exhibited potential disease-specific intercellular interactions involving Angptl4, Cxcl12, and Sema3 signaling. Disease-associated changes in these populations were confirmed by immunofluorescence in lung tissues from animals and patients.

**Conclusions:** We established a comprehensive cross-species single-cell atlas of mainstream rodent PH models, highlighting several novel macrophage and endothelial subtypes and signaling motifs potentially contributing to human disease.

## Introduction

Pulmonary arterial hypertension (PAH) is a heterogenous and progressive disease characterized by pulmonary vascular remodeling and increased pulmonary vascular resistance, frequently culminating in right heart failure.^1^ Endothelial cell (EC) dysfunction, smooth muscle cell (SMC) hyper-proliferation and hypertrophy, fibroblast-derived extracellular matrix and the infiltration of inflammatory cells contribute to pulmonary vascular remodeling in PAH, reflecting the involvement of multiple cell types in disease pathogenesis.^2–5^ While specialized subsets from these compartments may have unique roles in the pathogenesis of PAH, our current body of knowledge of the cellular mechanisms of PH has largely been limited to these general cell types. A comprehensive characterization of disease-associated resident pulmonary vascular and infiltrative cell populations and their signaling interactions at single-cell resolution would illuminate our understanding of the cellular mechanisms of PH.

An initial description of the single-cell transcriptomic profiles of patients with idiopathic pulmonary arterial hypertension (IPAH) demonstrated altered gene expression and signaling in pulmonary ECs, SMCs and fibroblasts.^6^ A subsequent single-cell study of rats exposed to monocrotaline (MCT) or SUGEN5416 combined with chronic hypoxia (SuHx) demonstrated differences in pulmonary and immune cells populations with model-specific profiles.^7^ Given the wide availability of genetically modified mice, murine models of PH have irreplaceable value in mechanistic studies of PH biology. Recently, a single-cell study focused on EC subtypes, revealing the roles of capillary EC populations in SuHx-exposed mice.^8^ The roles of other cell types and their interactions with ECs remain to be investigated. Importantly, widely used rodent models of PH, including exposure to chronic hypoxia (Hx), MCT or SuHx, are thought to exhibit overlapping but distinct histopathologic features reflecting different disease-causing mechanisms.^9^ Therefore, a cross-species analysis of these models at single cell resolution could help clarify their shared and distinct pathogenetic mechanisms.

In this study, we performed single-cell RNA sequencing (scRNA-seq) on the lungs of Hx mice, SuHx mice and MCT rats, and compared these to the human lung single cell atlas (HLCA) to define shared and unique perturbations in cell types, genes, and pathways across species and models. To ascertain mechanisms and features relevant to the pathogenesis of human PAH disease, these findings were in turn compared with available single-cell data from explanted human IPAH lungs,^6^ a large-scale bulk RNA profiling of PAH lung explants,^10^ and immunofluorescence staining of these cell subsets in experimental and human PAH lung tissues . We define roles of a novel macrophage subtype and functionally diverse EC subtypes that are shared and unique in these models, and signaling motifs within these subsets of potential relevance to human PH.

## Results

### Cross-species single-cell transcriptomic integration identified analogous immune cell populaions across models

PH phenotypes of hypoxia- and SuHx-treated mice and MCT-treated rats were confirmed by right ventricular systolic pressures and Fulton’s index (Figure S1). After quality control, a total of 68,078 cells were profiled in scRNA-seq datasets from these models (Figure S2). To integrate animal data with human reference data, single-cell transcriptomic data from these animal models were interpreted using the human lung cell atlas (HLCA) ^11^ based on homologous genes (Figure 1A-B). Utilizing the HLCA and canonical lineage marker genes, we identified 47 distinct PH-associated cell types, 38 of which were shared across all species (Figure 1C-E). To confirm analogous cell types, integrative non-negative matrix factorization was performed, and cell-type specific factors were identified. Correlations across species indicated that the general cell types from rodent models had high concordance with the corresponding populations in humans (Figure 1F, Figure S3). This approach to dataset integration has previously been validated, with high connectivity demonstrated between animal and human single-cell transcriptomes.^12^ In addition, disease-associated changes uncovered by animal single cell dataset were then validated in single cell transcriptomic data from human lungs affected by IPAH,^6^ and large-scale bulk transcriptomic data of human PAH lung explants ^10^ in subsequent analysis.

**Figure 1.**
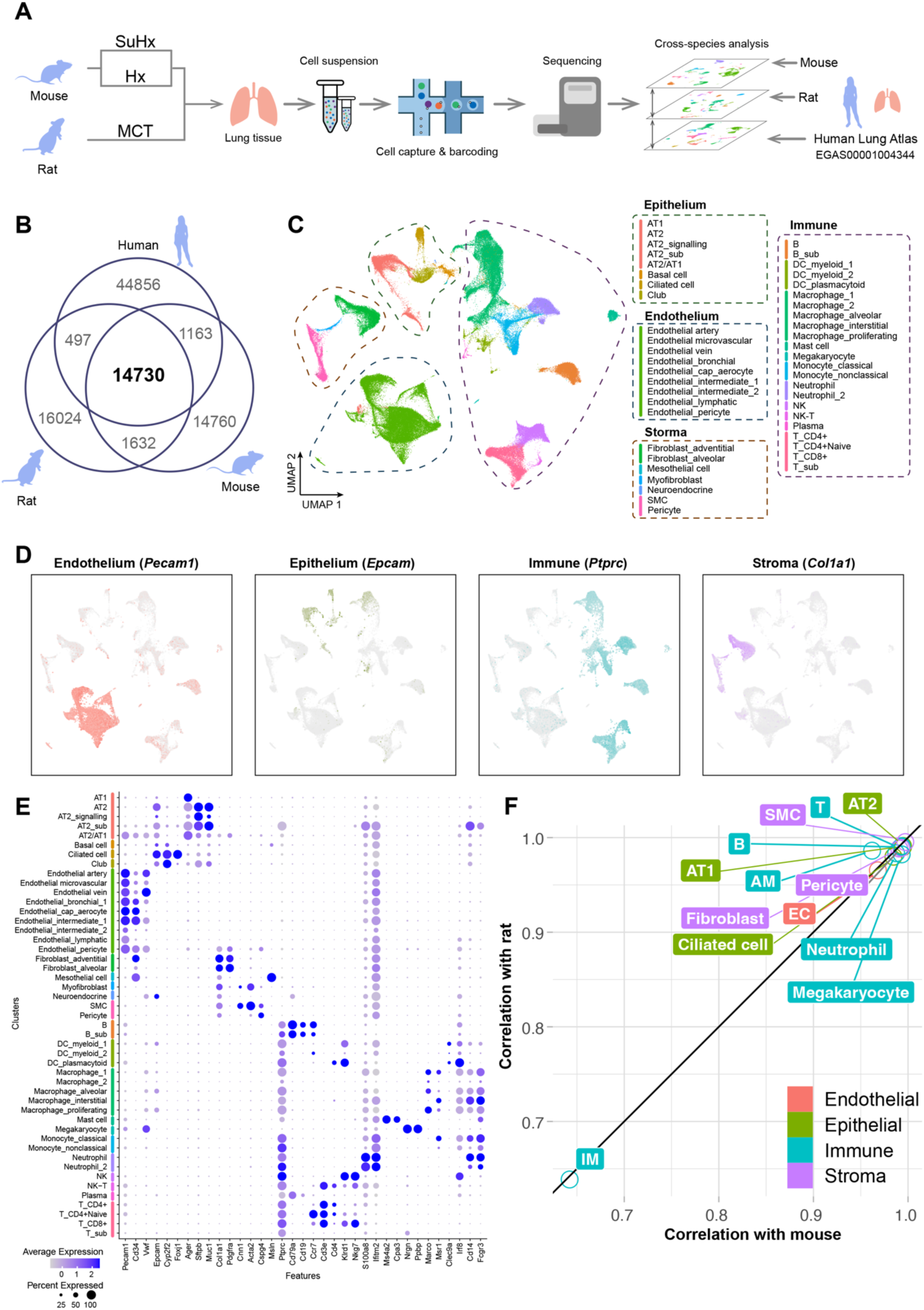
Integration of single cell transcriptomes from different models of pulmonary hypertension with human lung cell atlas reveals strong overlap in immune cell subsets. (A) Schematic diagram of study design for scRNA-seq on lung tissues from Hx mice, SuHx mice, MCT rats and control rats (Hx: n = 3, SuHx: n =3, MCT: n= 3, control mice: n = 5, control rats: n = 3). (B) Venn diagram depicts the numbers of homologous genes shared between species and thus available for cross-species integration. (C) Uniform manifold approximation and projection (UMAP) plot depicts lung cells from PH rodent models and human lung cell atlas with clusters labeled by cell type. (D) Classic markers for endothelial (*Pecam1*), epithelial (*Epcam*), immune (*Ptprc* encoding *Cd45*), and stromal cells (*Col1a1*) are superimposed on a cross-species UMAP. (E) Dot plot depicts gene expression levels and percentage of cells expressing canonical markers for each of the cell types using integrated data from all three species. (F) A correlation of cell types found in the human atlas versus rat and mouse atlases is shown. Pearson correlation coefficients were calculated between human, mouse, and rat using the dimensionally-reduced matrix generated by Non-Negative Matrix Factorization (NMF). Highly expressed genes (HEGs) for each cell type were calculated as the signatures, and cell types with fewer than 20 cells in each species were omitted from the analysis.

### Cell type prioritization emphasized the contribution of macrophages and endothelial cells

Cell type prioritization clarifies the contribution and/or changes of each cell type, evaluated by cell proportions, separability within cell types, transcriptomic changes, and thus potential relevance to a disease state.^13^ Based on cell proportions, microvascular endothelial cells (mECs) were significantly increased in SuHx-exposed mice but decreased in MCT rats. Dendritic cells and interstitial macrophages (IMs) were increased in MCT rats, while both neutrophils and macrophages were elevated in Hx-exposed mice (Figure 2A-B). The cell types perturbed in PH models were resolved and prioritized with respect to area under the curve (AUC) using the R package *Augur*,^13^ which quantified the separability of cells on a trained classifier. The results revealed that the most perturbed cell types were neutrophils and macrophages in Hx-exposed mice, ECs and macrophages in SuHx-exposed mice, and SMCs and macrophages in MCT rats (Figure 2C). To integrate transcriptomic changes, differentially expressed genes (DEGs) of each cell type were calculated for three PH models and combined into a union of DEGs (uDEGs) for each cell type, regarded as the transcriptomic changes of this cell type in PH. The mECs and IMs had larger numbers of uDEGs across all PH models (Figure 2D). The uDEGs of mECs were functionally enriched in endothelial migration, differentiation and angiogenesis, while the uDEGs of ΙΜs were enriched in the functions of T cell activation and leukocyte chemotaxis (Figure 2D). To assess the relevance of cell types to disease, 256 PH drug target genes were identified from the Cortellis Drug database (http://clarivate.com/products/cortellis/cortellis-competitive-intelligence). Using gene-set enrichment analysis (GSEA) of the highly expressed genes (HEGs), ECs, especially mECs, and macrophages were enriched with a greater proportion of PH-related genes (Figure 2E). Taken together, macrophages and ECs were the most modulated in frequency and expression profile, and thus appeared to be the most prominent contributing cell types in PH.

**Figure 2.**
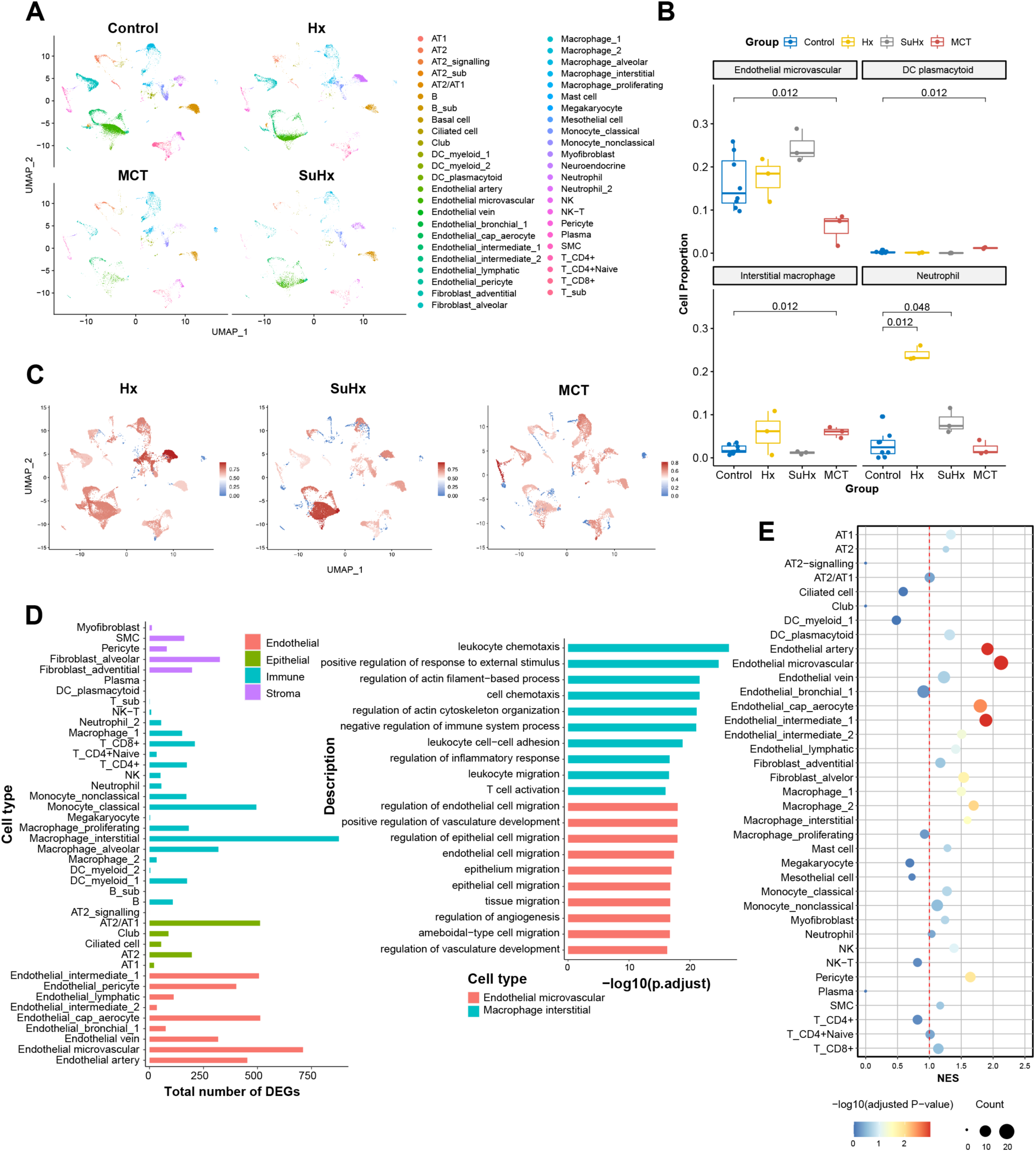
Cell type prioritization in response to perturbations in different PH models reveals the contribution of macrophages and endothelial cells. (A) UMAP plots depict lung cell types from control (mouse and rat), Hx (hypoxia-treated mouse), SuHx (SUGEN5416 and hypoxia treated mouse), MCT (monocrotaline-treated rat). (B) Box plots depict cell proportions of microvascular endothelial cells, dendritic cells, interstitial macrophages, and neutrophils from each model/species. Statistical significance was tested by Wilcoxon rank sum test. (C) The area under curve (AUC) calculated by Augur is depicted by heat map color on UMAP plots reflecting cell prioritization from multiple experimental PH models. (D) Bar plots show the union of differentially expressed gene sets (uDEGs) from multiple PH models (left panel) and gene set enrichment analysis (GSEA) of uDEGs from microvascular endothelial cells and interstitial macrophages (right panel). Differentially expressed genes of each cell type were calculated by the Seurat FindMarker function in each of the different PH models. The uDEGs of one cell type was defined as the combined set of DEGs from all three models, and was considered in subsequent analyses to be the PH-specific transcriptomic signature for this cell type. (E) Dot plot shows the normalized enrichment score (NES) of PH drug target genes in various cell types, determined by GSEA of highly expressed genes (HEGs) from each cell type with the Cortellis Drug Discovery Intelligence platform.

### Macrophage subtype analysis revealed pro-inflammatory polarization of alveolar macrophages

To investigate their heterogeneity further, macrophages were re-clustered, and GSEA was performed on the HEGs of macrophage subtypes to assess their enriched functions. Considering the concordance with human IPAH, cell subtype analyses were performed on the single-cell dataset from human IPAH to validate key findings in the animal single-cell data. Macrophage subtypes were annotated as previously described ^14,15^ using four primary functional subtypes: tissue-remodeling, pro-inflammatory, cytokine-producing, and proliferative (Figure 3A-C). Tissue-remodeling macrophages were enriched in the functions of wound healing, cell-substrate adhesion and angiogenesis (Figure 3C), potentially representing an M2-like profile ^16^. The transcriptome of tissue-remodeling macrophages resembled the *FABP4*^+^ macrophages previously described as the main alveolar macrophage (AM) subtype in bronchoalveolar lavage fluid.^17^ IMs were annotated as pro-inflammatory due to the enriched functions of leukocyte differentiation and migration, and the production of pro-inflammatory factors IL1β, IL6 and TNFα (Figure 3C), which were M1-like and consistent with the reported transcriptomic signature of IMs.^18^ On the other hand, cytokine-producing macrophages highly expressed genes involved in cytokine production and regulation of cell proliferation, such as *Prg4* and *Igf1* (Figures 3C-D).

**Figure 3.**
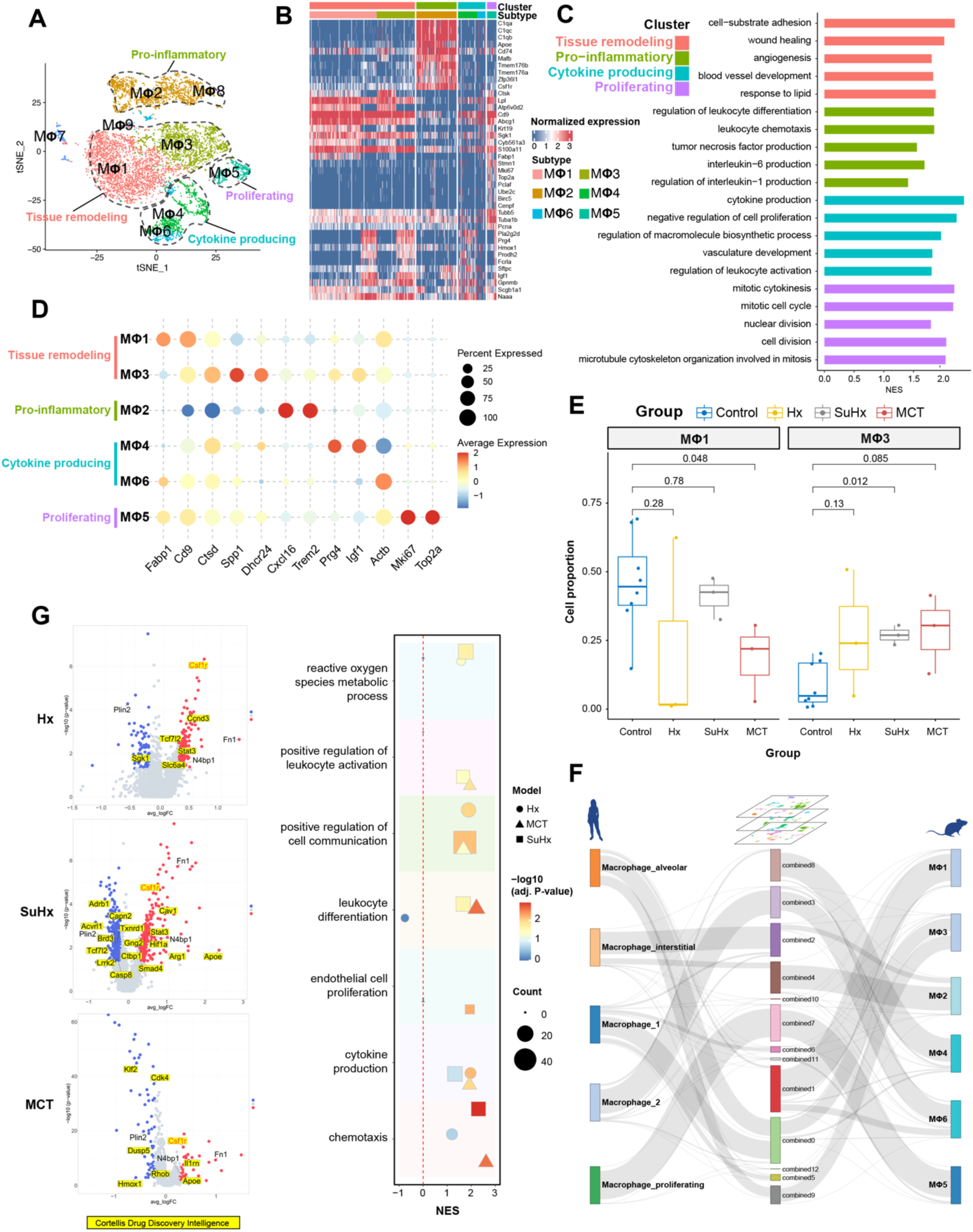
Analysis of macrophage heterogeneity across multiple PH models reveals pro-inflammatory polarization of an alveolar macrophage subset. (A) A t-distributed stochastic neighbor embedding (t-SNE) projection of macrophages from Hx, SuHx, MCT, and control groups is shown with circled regions reflecting the functional annotation of distinct macrophage subtypes. (B) A heatmap of the top 10 HEGs for each of the macrophage subtypes is shown, ranked by log fold change. (C) Summary of GSEA comparing each of the macrophage subtypes. (D) Dot plot shows the expression of key genes in each of the macrophage subtypes. (E) Box plot shows cell proportions of two clusters of tissue remodeling macrophages, with statistical significance determined by Wilcoxon rank test. (F) DEGs within the M<λ3 macrophage subtype are depicted by volcano plot with respect to each of the PH models/species (left panel), with a summary of GSEA corresponding to these DEGs (right panel). DEGs overlapping with PH drug target genes from Cortellis Drug Discovery Intelligence are highlighted, and one consistently upregulated gene, *Csf1r*, is labeled in red. (G) A river plot shows connectivity and similarity of macrophages between human and animal datasets.

Tissue-remodeling macrophages could be further sub-divided into two distinct clusters (Figure 3A), MΦ1 was the main subtype in control animals, whereas MΦ3 was increased in all PH models (Figure 3E). While retaining the M2-like signature of tissue-remodeling macrophages, MΦ3 highly expressed pro-inflammatory genes (e.g. *Il1b*, *Spp1*, *Ccl2*) and genes involved in sterol biosynthetic and metabolic processes (e.g. *Sqle*, *Dhcr24*) as compared with MΦ1 (Figure 3D and Figure S4A). MΦ3 exhibited a hybrid signature of both M1 and M2 macrophage subsets, and thus appeared analogous to a previously identified *SPP1*^hi^ AM population in IPAH ^6^ (Figure S5A-B) and IPF patients ^17^. Based upon being a PH target gene within the top 10 HEGs in this population, we selected *Dhcr24* as a marker for the MΦ3 subpopulation, a.k.a., *Dhcr24^high^* tissue-remodeling macrophages. For cross-species analysis, a highly analogous *DHCR24^high^*human alveolar macrophage cluster was mapped and projected onto various clusters from the rat and mouse PH models and subjected to UMAP analysis. (Figure 3F, Figure S4B). Consistent with a strong correlation between PH models and human disease, *DHCR24* was upregulated in AMs from IPAH patients, and there was a trend of an increased proportion of *DHCR24^high^* AMs.^6^ (Figure S5C-D).

DEGs of the *Dhcr24^high^* tissue-remodeling macrophages included numerous PH target genes, suggesting a key role in PH. In particular, *Csf1r* (macrophage colony stimulating factor 1 receptor) was upregulated in this cluster across all models, as well as in IPAH patients (Figure 3G, Figure S5E).^6^ DEG enrichment revealed shared and unique functional changes in the *Dhcr24^high^*tissue-remodeling macrophage cluster across the three models. In the SuHx and MCT models, GSEA revealed upregulated genes of MΦ3 *Dhcr24^high^* tissue-remodeling macrophages to be related to leukocyte chemotaxis, activation and differentiation, whereas in SuHx-exposed mice, genes relevant to EC proliferation were upregulated (Figure 3G, right panel), suggesting that these macrophages might contribute to the dysregulated endothelial angiogenesis and angio-obliterative phenotype in this model. In the Hx and MCT models, cytokine and reactive oxygen species (ROS) production were upregulated (Figure 3G). Across all three models, *Dhcr24^high^*tissue-remodeling macrophages showed upregulated cell communication functions that could reflect a mediator role via interactions with other cell types driving PH (Figure 3F). Taken together, this specialized population appears to have an important role in the development of PH across models and species, but with slightly different mechanistic contributions highlighted in each model.

Via immunofluorescent staining, we found an increased abundance of Dhcr24^+^Csf1r^+^Cd68^+^ macrophages corresponding to MΦ3 *Dhcr24^high^* tissue-remodeling macrophages in the mouse (Figure 4A) and rat PH models vs. controls (Figure 4B), as well an increased abundance of DHCR24^+^CSF1R^+^CD68^+^ in IPAH patient-derived lung tissues vs. control lungs (Figure 4C), Moreover, compared with non-diseased control samples, the ratio of DHCR24^+^CSF1R^+^ cells among total CD68^+^ macrophages was significantly increased in all models and in human PAH tissues (Figure 4D). The ratio of DHCR24^+^CSF1R^+^ cells to total CD68^+^ macrophages was significantly increased in Hx mouse model (p=0.0484, 0.053±0.020 vs. 0.21±0.033), the SuHx mouse model (p=0.0002, 0.053±0.020 vs 0.38±0.064), in MCT rats (p<0.0001, 0.057±0.0070 vs. 0.36±0.036), and human PAH tissues (p=0.031, 0.065±0.013 vs. 0.24±0.053; n=6 per group, data shown as mean ± SEM, One-way ANOVA, Dunnet’s test for multiple comparisons).

**Figure 4.**
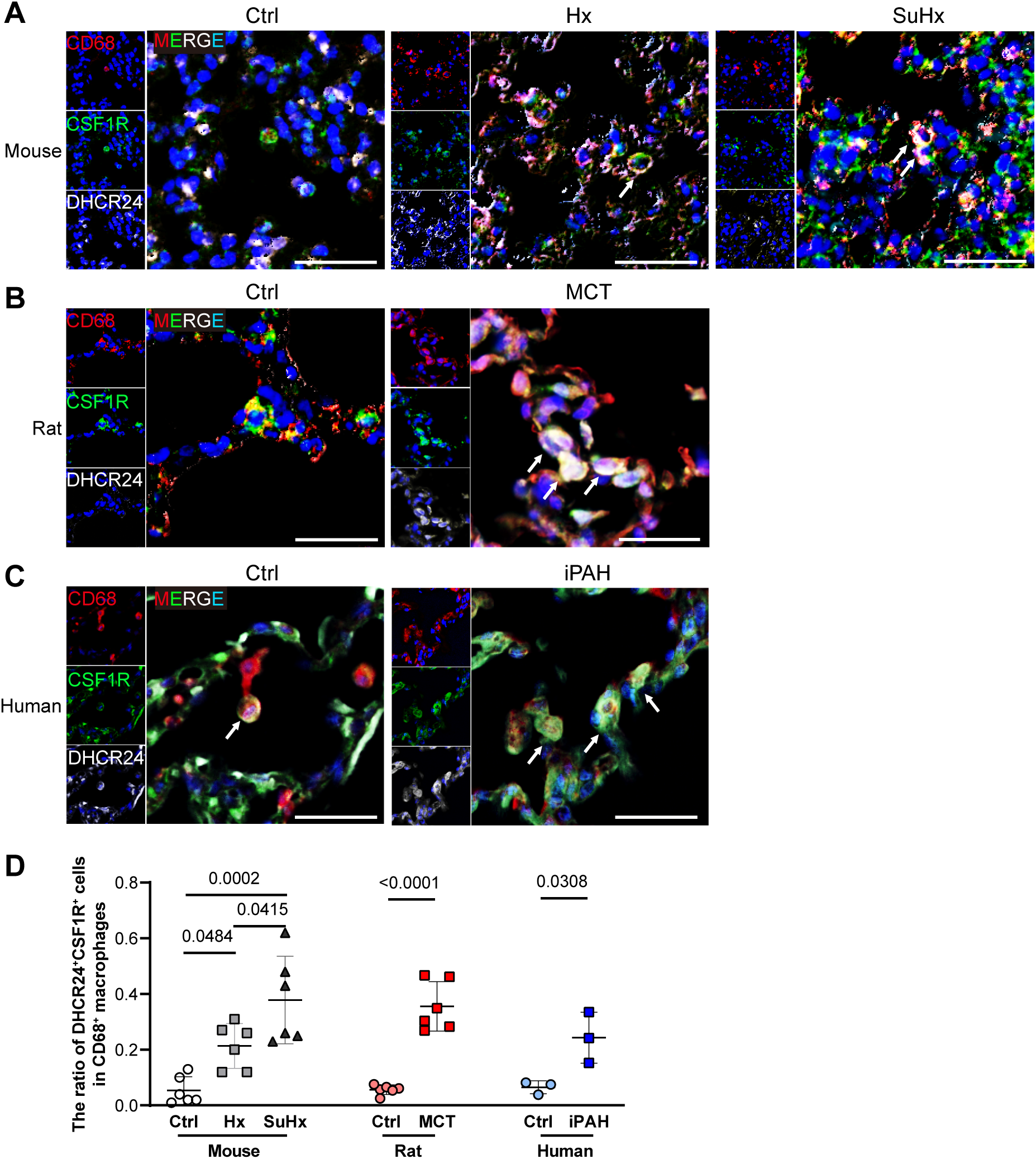
*Dhcr24*^high^ macrophages are enriched in experimental PH and human PAH tissues by immunofluorescent staining. Macrophages from the M<λ3 subtype were defined by the expression of a pan-macrophage marker (CD68^+^, red) in combination with DHCR24 (white) and CSF1R (green). These M<λ3 macrophages were enriched in mouse PH vs. controls **(A)**, and rat PH vs. controls **(B)**, and human IPAH tissues versus controls **(C)**. Cell nuclei were counterstained with DAPI (blue). White arrows indicate the DHCR24^+^CSF1R^+^CD68^+^ cells. Scale bars=25μm. (**D**) Quantification of immunofluorescent staining, showing the consistent increase of DHCR24^+^CSF1R^+^CD68^+^ macrophages in rodent models of PH (n=6 per group) and iPAH patients (n=3 per group). One-way ANOVA was used for comparison in the mouse models, and t-test was used in the rat model and human tissues. Data are presented as the mean ± SEM.

Given the increased abundance of *Dhcr24^high^* macrophages in diseased animals and patients, and the reciprocal decrease in the MΦ1 population, we postulated that common cell populations transition between these two closely-related subtypes of tissue-remodeling macrophages. The results of monocle trajectory analysis ^19,20^ indicated a transitional trend between these macrophage subtypes (Figure S6A-D). The pseudo-time and DEGs from monocle were utilized to establish a least absolute shrinkage and selection operator (LASSO) regression to identify key genes involved in this transition (Figure S6C-D). GSEA of the key genes showed an upregulation of IL-6 and TNFα production, and cellular responses to cytokines, and a downregulation of lipid modification and TGFβ signaling (Figure S6D, right panel), reminiscent of the polarization between M1 and M2 macrophages. Based on the pseudo-timeline and pseudo-time distribution of the two clusters in the Hx and MCT models, tissue remodeling macrophages may polarize towards *Dhcr24^high^* cells and acquire pro-inflammatory features in these disease models. In SuHx-exposed mice, *Dhcr24^high^* macrophages potentially polarized towards MΦ1 to promote EC proliferation and vascular remodeling (Figure 3E, Figure S6E). Additionally, 10 transition-related key genes were found in the PH drug target database, indicating that the transition between macrophage subtypes may contribute to PH pathogenesis.

### Heterogeneity of endothelial cells revealed functionally distinct subtypes corresponding to model-specific pathological changes

After re-clustering ECs, highly expressed genes of each EC clusters were calculated and used for GSEA enrichment analysis. EC subtypes were annotated based on the enriched functions known to be associated with PH. We focused on the microvascular ECs (mECs) that were further annotated into 6 functionally diverse subtypes and highly heterogenous between the experimental groups (Figure 5A-B). Normal ECs were the main population of mECs in the control group with high expression of *Plvap* (Figure 5C), a signature gene for mECs ^21,22^. By contrast, apoptotic ECs (*i.e.,*EC2, EC4, EC16) were enriched in GSEA for functions of apoptotic process and cell chemotaxis, and were significantly increased in the Hx and SuHx models (Figure 5B). Consistent with pulmonary endothelial populations in the HLCA, four clusters (EC5, EC6, EC12, EC13) were highly enriched for genes characteristic of the pulmonary vasculature with high expression of *Endrb*, *Kdr*, and *Hopx*, analogous to the previously identified aerocyte capillary (aCap) and general capillary (gCap) populations ^11^ (Figure 5B-C). Two of these clusters (EC5, EC6) showed signatures of aerocyte ECs, lacked several classic EC markers, and were enriched in blood vessel endothelial migration function (Figure 5B-C). By contrast, EC12 had mixed features of normal mECs and aerocyte ECs, and were enriched in muscle cell differentiation but not in endothelial migration (Figure 5B). *Ednrb,* which encodes endothelin receptor type B, was highly expressed in EC5, EC6 and EC12 clusters and distinguished these from other EC subtypes as one of top 10 HEGs. We therefore annotated EC5 and EC6 as *Ednrb*^+^ aerocyte ECs (aECs) and EC12 as *Ednrb*^+^ vascular ECs (vECs). Additionally, EC13 were more similar to apoptotic ECs (Figure S7). Notably, *Ednrb*^+^ vECs mainly existed in the control group but were lost in all three PH models (Figure 5D). Consistent with its diminished expression in experimental PH, expression of *EDNRB* mRNA was decreased in total lung tissues from PAH patients.^10^(Figure S8A). Additionally, *EDNRB^+^* ECs were detected in the single-cell dataset of human IPAH but could not be subdivided further due to limited sequencing depth and cell numbers (Figure S8A). Moreover, an increased abundance of EC11, named as angiogenic EC, was seen almost exclusively in SuHx-exposed mice (Figure 5D), with a signature enriched by GSEA for angio-obliterative functions (*i.e.,* cell proliferation and angiogenesis, Figure 5B), possibly corresponding to aberrantly proliferating or apoptosis-resistant EC subtypes in the SuHx mice and plexiform lesions in PH patients. EC17 cells, marked as *Nox2^+^* ECs, were increased in MCT rats and rarely observed in any other models (Figure 5A-B and D), with highly expressed genes related to ROS metabolism and inflammatory responses, signifying a newly identified model-specific EC subtype.

**Figure 5.**
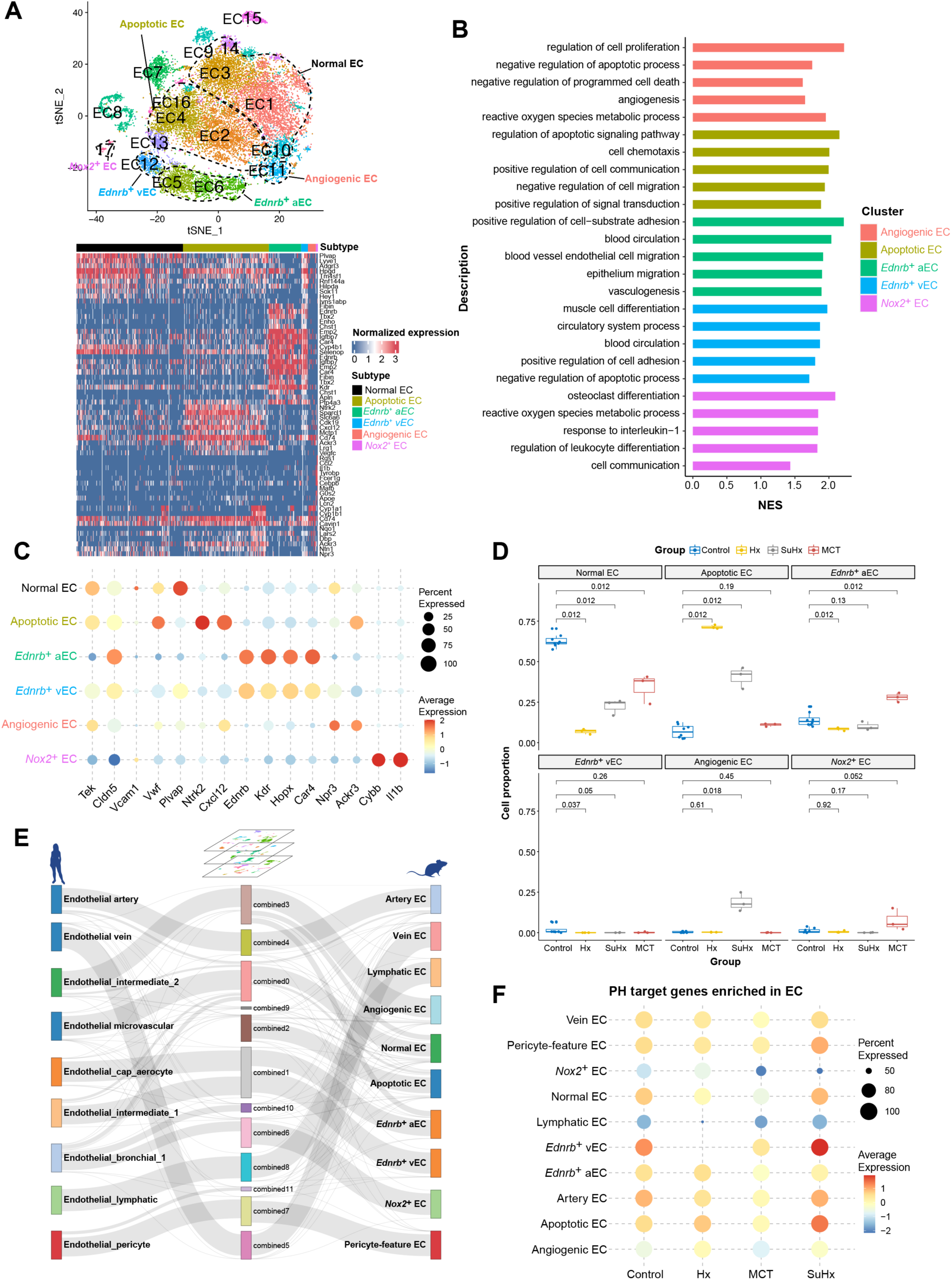
Analysis of endothelial cell heterogeneity reveals functional EC subtypes corresponding to PH disease across species and models. (A) A t-distributed stochastic neighbor embedding (t-SNE) projection of ECs from Hx, SuHx, MCT, and control groups is shown with circled regions reflecting the functional annotation of distinct EC subtypes, including the Ednrb+ microvascular subset (EC12 cluster, upper panel). A heatmap of the top 10 HEGs for each of the EC subtypes is shown, ranked by log fold change (lower panel). (B) Summary of GSEA comparing microvascular EC subtypes. (C) Dot plot showing the expression of key genes in each of the EC subtypes. (D) Box plot showing cell proportions of microvascular EC subtypes, with statistical significance tested by Wilcoxon rank test. (E) River plot reveals connectivity and similarity of EC subsets between species and models. (F) Dot plot depicts the expression of PH target genes enriched in ECs.

In the cross-species analysis, analogous human mECs subpopulations could be matched with each of these animal models (Figure 5E and Figure S8B-C). Since PH target genes were most enriched in mECs (Figure 2E), each subtype was scored by these genes (Figure 5F). *Ednrb*^+^ vECs and apoptotic ECs showed more relevance to PH, further indicating an important role of these subtypes. Moreover, analogous clusters for each of the EC subtypes (*i.e.* apoptotic, angiogenic, *EDNRB^+^* and *NOX2^+^* ECs) were found in iPAH patients ^6^ (Figure S8D-E). In addition to mECs, EC9 corresponded to ECs from pulmonary arteries characterized in a previous study in human PAH.^23^ Taken together, these data suggested that while ECs exhibited substantial heterogeneity across multiple models, and there was convergence of several functional EC subtypes reflective of pathogenetic cell populations in human disease, and divergence of several clusters that may reflect the unique pathophysiology of each experimental model.

To validate the functionally diverse EC subtypes, we selected marker genes based on their functions and specificity. Among the mECs expressing PLVAP, *Ednrb*^+^ vECs were found to co-express ETB (endothelin receptor type B encoded by *Ednrb*), KDR and HOPX. The results of immunostaining demonstrated that ETB^+^ vECs were prominent in the endothelium of pulmonary micro vessels (<50 μm diameter) in control lungs but were markedly reduced in the remodeled pulmonary vessels in PH models and iPAH patients (Figure 6A). The ratio of ETB^+^ vECs to total mECs was significantly decreased in Hx mouse model (*p*<0.0001, 0.38±0.039 vs. 0.017±0.0057), the SuHx mouse model (*p*<0.0001, 0.38±0.039 vs 0.082±0.005), in MCT rats (*p*<0.0001, 0.57±0.033 vs. 0.015±0.0034), and human PAH tissues (*p*=0.0001, 0.48±0.031 vs0.022±0.0016; n=6 per group for experimental models, n=3 per group for human tissues, data shown as mean ± SEM, One-way ANOVA, Dunnet’s test for multiple comparisons) (Figure 6B). Apoptotic ECs were defined as NTRK2^+^. *Ntrk2* was one of the top HEGs among endothelial subpopulations, encoding a neurotrophic tyrosine receptor kinase. These cells were readily visualized in the endothelium of PH mouse models and iPAH patients (Figure 6C). The ratio of NTRK2^+^ vECs to total mECs was significantly increased in Hx mouse model (*p*=0.0028, 0.052±0.012 vs. 0.25±0.059), the SuHx mouse model (*p*<0.0001, 0.052±0.012 vs 0.50±0.014), and human PAH tissues (*p*=0.0006, 0.037±0.031 vs0.24±0.019). For rat-specific ECs, NOX2 (encoded by *Cybb*) was selected owing to its specificity and importance for ROS-mediated interactions between inflammatory and endothelial cells during vascular inflammation.^24^ Immunostaining showed significantly increased NOX2^+^ ECs in the endothelium of MCT rats (*p*=0.017, 0.066±0.014 vs. 0.17±0.034) and human PAH tissues (*p*=0.030, 0.024±0.0075 vs0.23±0.061), suggesting the presence of ROS-mediated damage (Figure 6E-F).

**Figure 6.**
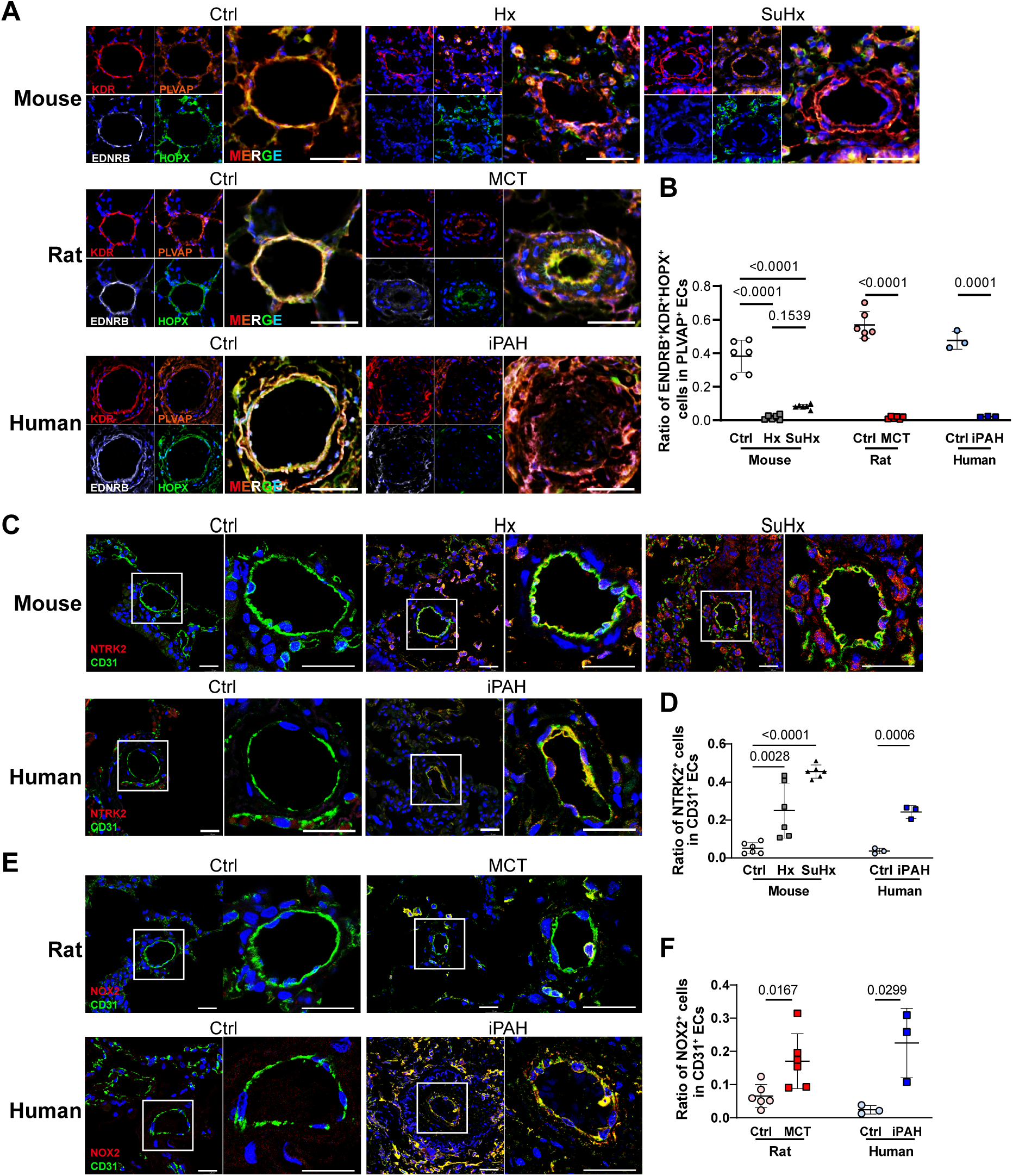
Specific EC subtypes that are perturbed across experimental PH and human PAH or unique to individual models by immunofluorescent staining. (**A**) *Ednrb*^+^ vECs (cluster 12) were marked with EDNRB (white), KDR (red), HOPX (green) and PLVAP (orange) and representative images were showed in the lung sections of Ctrl vs. Hx and SuHx mice, Ctrl vs. MCT rats, or control subjects vs. iPAH patients. (**B**) Quantification of immunofluorescent staining, showing the consistent decrease of EDNRB^+^KDR^+^HOPX^+^PLVAP^+^ vECs across rodent models of PH and iPAH patients. (**C**) Apoptotic ECs (cluster 4) characterized as NTRK2 (red) and CD31(green) double positive cells were most prominently enriched in hypoxia-treated mice. (**D**) Quantitative immunofluorescence reveals increased abundance of apoptotic ECs across mouse models of PH and iPAH patients. (**E**) *Nox2*^+^ ECs were determined using NOX2 (red) and CD31(green) antibodies and representative results in the lung sections of rats and human were shown. (**F**) Quantitative immunofluorescent staining reveals increased frequency of NOX2^+^ ECs in the MCT model and iPAH patients. Cell nuclei are counterstained with DAPI (blue). White arrows indicate the positive immunostaining in different groups. Scale bars = 25μm. One-way ANOVA was used for comparison in the mouse models, and t-test was used in the rat model and human tissues. Data are presented as the mean ± standard error.

### Macrophages and endothelial cells showed regulatory cell communications

Given the striking changes in the macrophage and endothelial subpopulations associated with PH, we investigated the intercellular communication between these highly perturbed cell types using *CellChat*.^25^ *Ednrb*^+^ vECs had a high probability of interactions with normal ECs, tissue-remodeling macrophages, and themselves (Figure 7A). Specifically, *Ednrb*^+^ vECs were predominant source cells in apelin and c-KIT pathways that interacted with Normal ECs, indicating a protective role of *Ednrb*^+^ vECs against PH (Figure 7B). Conversely, apoptotic ECs, angiogenic ECs, and normal ECs might modulate *Ednrb*^+^ vECs by semaphorin 3 signaling (Figure 7B). Angiopoietin-like signaling was abundant among EC subtypes, to which the tissue-remodeling macrophages were the main source of the ligand *Angptl4* (Figure S9A). The expression of *Angptl4* was reduced in all PH models and iPAH patients,^6^ probably resulting from changes in expression associated with polarization from *Angptl4^high^* MΦ1 to *Angptl4^low^* MΦ3 (Figure S10A-C). Importantly, the probability of cell interactions varied in the three models (Figure 7C). In Hx-exposed mice, *Cxcl12* secreted by normal and apoptotic ECs showed an increased interaction with *Cxcr4* on pro-inflammatory and tissue-remodeling macrophages (Figure 7D, Figure S9A). The expression of *CXCL12* was increased in the microarray data of PAH patients ^10^ (Figure S10D), which was consistent with the results from PAECs isolated from IPAH patients. In SuHx-exposed mice, semaphorin 3 signaling increased between apoptotic ECs and angiogenic ECs, which might be related to the dysregulated angiogenic signaling in this model ^26,27^ (Figure 7E). The activation and increase of SEMA3 pathway was consistent with the findings in the recent single cell study of pulmonary arteries ^28^. In MCT rats, pro-inflammatory and tissue-remodeling macrophages secreted more *Hbegf*, potentially contributing to mEC dysfunction (Figure 7F, Figure S9C). Additionally, IGF signaling was more abundant in *Nox2*^+^ ECs with ligands secreted by macrophages (Figure 7F). Furthermore, an apparent increase in autocrine EPHB signaling of apoptotic ECs was found in the Hx and SuHx models (Figure 7G). Taken together, cell communications related to endothelial function and macrophage chemotaxis were abundant in PH, but the distinct communicating cells might be related to the model-specific pathologic changes.

**Figure 7.**
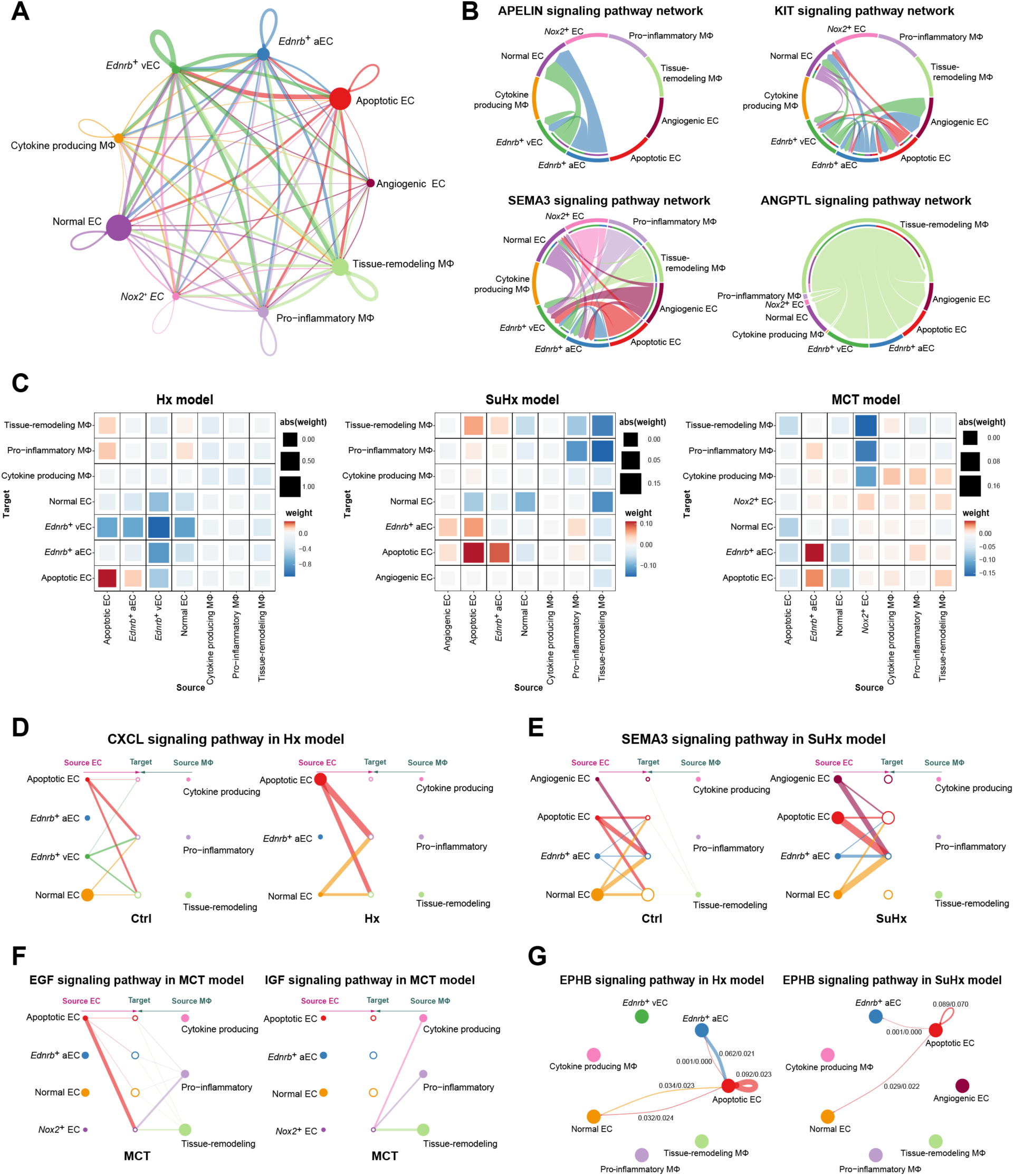
Cell communication between macrophages and endothelial cells. (A) Cellchat analysis reveals significant ligand-receptor pairs between cell populations, with edge width proportional to the number of ligand-receptor pairs. (B) Chord diagram showing apelin (APELIN), c-Kit (KIT), semaphorin (SEMA3), and angiopoietin-lke (ANGPTL) signaling pathways among macrophage and EC subtypes. (C) Heatmap reflects changes in the weight of significant ligand-receptor pairs between any pair of two cell populations with respect to each of the three PH models. (D-F) Hierarchical plots show perturbed signaling pathways in each of the three PH models: Cxcl12 (CXCL) in the Hx model; semaphorin 3 (SEMA3) in the SuHx model, and EGF/IGF in the MCT model. For D and E, left panels show the signaling pathway in control group, and right panels show changes in PH models. In the MCT model, there is no significant interaction of EGF and IGF in the control group. (G) Circle plot showing the changes of weight of ephrin B (EPHB) signaling pathway in the Hx and SuHx models, showing increased autocrine signaling of apoptotic ECs.

## Discussion

This is the first study comparing the single-cell atlas of mouse and rat models of PH combined with the HLCA and human PAH transcriptomic and single cell transcriptomic datasets, uncovering shared and unique mechanisms between animal models and human PAH at single-cell resolution. The findings in animal models were mapped to drug targets in patients, revealing enrichment of drug target genes in macrophages and mECs. A newly identified DHCR24^high^ subtype was consistently increased among macrophages in PH models and human patients with reciprocal decreases in an MΦ1 subset to suggest increased polarization with disease. Functionally diverse mEC subtypes were identified and closely related to the model-specific pathological phenotypes, of which EDNRB+, pro-angiogenic and NOX2+ subpopulations were prominent and shared with IPAH. Cell communications between macrophages and ECs highlighted signaling motifs known to be important in PH, including CXCL12, Apelin, EPH, and Hbegf axes previously valdated in PAH, and revealed several novel pathways including ANGPTL4 and SEMA3 (Figure 8).

**Figure 8.**
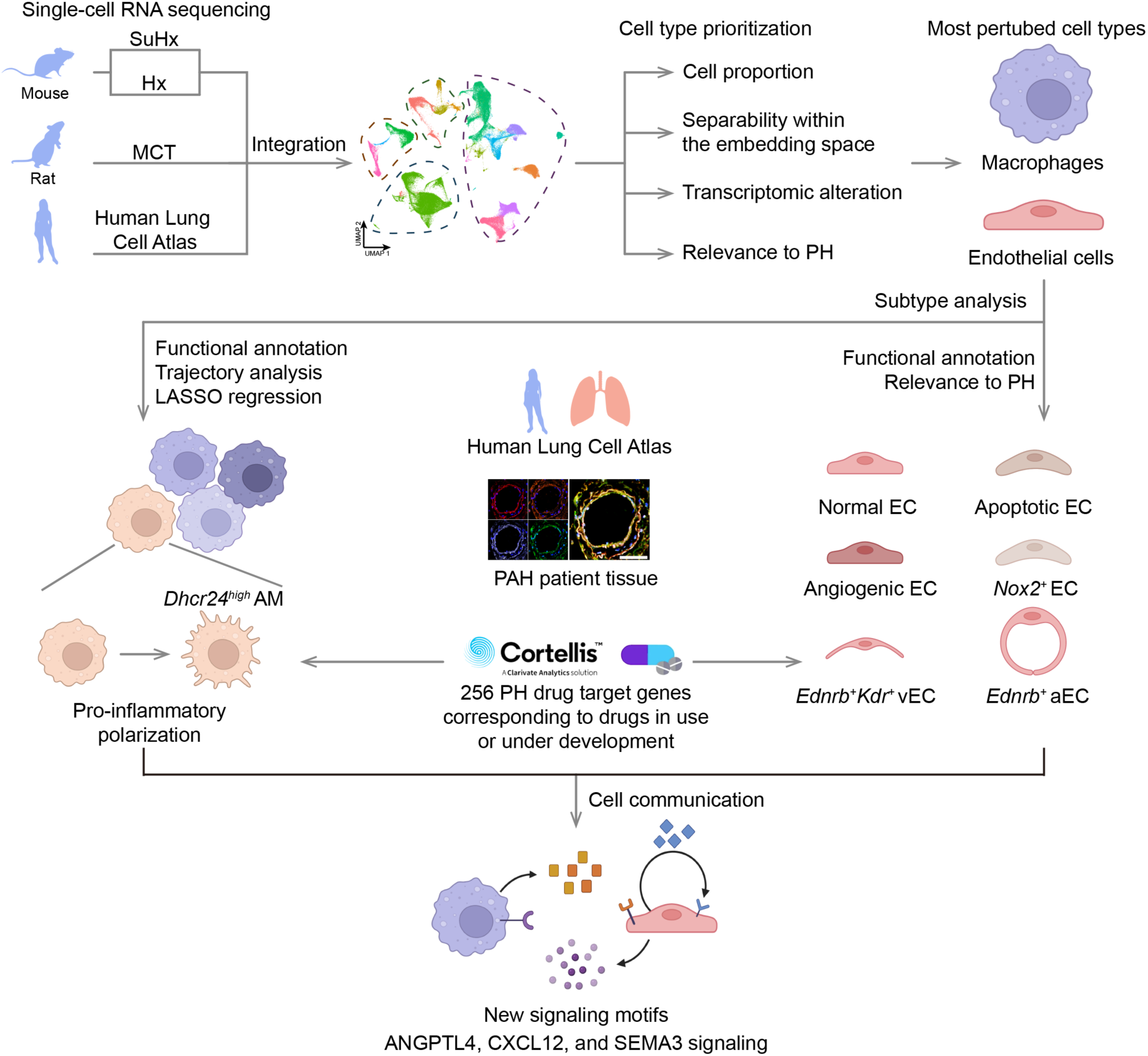
Schematic diagram of the workflow in this study. ScRNA-seq was performed on mouse and rat models of PH and the data were integrated with the human lung cell atlas. By assessing cell type prioritization, macrophages and endothelial cells were found to be the most perturbed cell types in PH and were further separated into subtypes. Novel cell subtypes, including *Dhcr24^high^* AMs and *Ednrb*^+^ vEC, were identified to be perturbed in experimental PH, and confirmed in PAH patient tissues. These new macrophage and endothelial subsets had transcriptional signatures that overlapped highly with known PH drug target genes, and demonstrated evidence of intercellular communication consistent with both established and novel signaling motifs for PH.

Rodent models do not fully recapitulate all characteristics of human PH but replicate many key features. Commonly used Hx and SuHx murine models do not develop severe pathological changes,^29^ but mice are indispensable in mechanistic investigations owing to their genetic maneuverability. SuHx exposure causes angio-obliterative remodeling in rats, resembling the plexiform lesions in patients.^29,30^ In a seminal study, single-cell transcriptomic analysis MCT- or SuHx-exposed rats identified molecular targets for drug repurposing.^7^ Our data were complementary to this report, but have incorporated two widely-used mouse models in addition. Consistent with the prior report, we found evidence of increased DCs and IMs in MCT rats. Importantly, our current study analzyed larger numbers of cells and performed sequencing with greater depth to reach a saturated number of genes (median of detected genes: 1699), yielding the most comprehensive analysis of cell subtypes in experimental PH to date. Taken together, we have produced a comprehensive cross-species dataset for translational studies and identified functionally distinct cell populations unique to specific models, supporting their utility for investigating specific aspects of human PH.

Macrophages are increasingly recognized for their importance in PH. Depletion of AMs was previously found to attenuate hypoxia-induced PH.^31^ We annotated two AM subtypes and found an increase of *Dhcr24^high^* tissue-remodeling macrophages in diseased animals, consistent with published data from SuHx rats (Figure S11A-C). The expression of *Dhcr24* was not limited to macrophages but distinguished the two subtypes of tissue-remodeling macrophages. Our data predicted the transition of AMs towards a pro-inflammatory phenotype in PH. In agreement with the reported activation of AMs into a hypoxic program, *Dhcr24^high^*tissue-remodeling macrophages represented most of AMs, and showed upregulated mTOR and EIF3m/EIF4ebp1 Figure S11D), pathways that were previously reported to be critical for the activation of AMs ^32^. The genes involved in AM transition included known PH targets, such as *Gsr* (macrophage polarization),^33^ *Sphk1* (macrophage activation),^34^ and *Nampt* (differentiation and polarization).^35^ These results supported the importance of AM transition in PH pathogenesis. For IMs, an increase in both the Hx and MCT models and the high expression of chemotaxis-related genes are consistent with the expansion of inflammatory macrophages driven by monocyte chemotaxis, contributing to PH.^36^ Increased thrombospondin-1 derived from IMs contributed to hypoxia-induced PH.^37^ This change was observed in Hx-exposed mice (Figure S11E), but not in other models, indicating their heterogeneous cellular mechanisms. In recent single cell studies, Kumar et al. focused on the subtypes of IMs exposed to acute and prolonged hypoxia and demonstrated dynamic changes of inflammatory regulation during hypoxia exposure^38^. Taken together, in contrast to traditional strategies classifying macrophages into M1/M2 or alveolar/interstitial types, our single-cell analysis provides an extensive characterization of macrophage subtypes, and postulates the transition between these subtypes in PH pathogenesis.

ECs are known to be important in PH but have been generally studied as a single cell type. Building upon the EC subtypes reported in the landmark single-cell studies of PH,^8,23^ this study has refined the characterization of ECs to an unprecedented depth, using 21,564 ECs to classify functionally distinct populations contributing to disease pathogenesis. We identified EC subtypes corresponding to key pathological features or signaling pathways, such as angiogenic ECs in SuHx-exposed mice and *Ednrb*^+^ vEC in the control group, and confirmed our prediction by experiments. This ETB-expressing EC subtype might have potentially protective roles by mediating vasodilation.^39,40^ Mutations of *KDR* (a.k.a. *VEGFR2*) are found in patients with PAH,^41,42^ and *Kdr* blockade is related to aberrant endothelial proliferation and PH,^43,44^ suggesting an important role of these *Ednrb*^+^ vECs co-expressing *Kdr*, which were identified in this study for the first time*. Hopx* was also enriched in this EC subtype, but its role deserves further investigation. Additionally, apoptotic ECs with high *Ntrk2* expression were the major EC subtype in PH models and increased in patients. As *Ntrk2* kinase participates in inflammation and apoptosis,^45,46^ the data revealed *Ntrk2* as a potential target against EC apoptosis. Furthermore, *Nox2*^+^ ECs were associated with ROS and inflammation, while their roles in PH remain unclear despite previous studies.^47,48^ By scRNA-seq, we found this *Nox2*^+^EC subtype in MCT-exposed rats, which could represent an adequate model for investigating EC-related ROS. Collectively, we have defined ECs into novel subtypes with different transcriptomic profiles and functions suggestive of protective or detrimental roles and elucidated the unique pathological features of the different models, facilitating model selection in future investigations.

By inferential analysis of cell communication, we identified potential ANGPTL signaling between tissue-remodeling macrophages and all EC subtypes, with macrophages being the main source of *Angptl4*, a regulator of vascular homeostasis and angiogenesis.^49,50^ CXCL12 signaling was found to be crucial in PH as it regulates the infiltration of macrophages ^51^ and the migration and proliferation of SMCs.^51,52^ Consistent with this notion, we found increased CXCL12 signaling in ECs of Hx-exposed mice, with receptor *Cxcr4* expressed on macrophages and SMCs (data not shown). Importantly, *Cxcl12* was most upregulated in apoptotic ECs, which were the principal cellular source of this ligand. Furthermore, we found possible cell communications among EC subtypes, SEMA3 signaling in SuHx-exposed mice may promote aberrant angiogenesis,^53,54^ as SEMA3 ligands were secreted by dysfunctional ECs and acted on the protective *Ednrb*^+^ ECs. Intriguingly, apoptotic ECs demonstrated autocrine EPHB signaling related to vasculogenesis and angiogenesis,^55^ potentially giving rise to hyper-proliferative cells. Taken together, the findings allowed us to predict key signaling motifs, illustrating the source of signals and the target cell populations, providing new insight on the cellular interactions resulting in pathological vascular remodeling.

We determined the cell type-specific expression of drug target genes. As expected, these target genes were more abundant in ECs. Among the endothelin receptor antagonists currently used for PAH, ambrisentan and macitentan more selectively target the ETA receptor, which is located on smooth muscle cells and may have opposing effects to the ETB receptor.^40,56^ We found *Ednrb* to be specifically expressed on certain EC subtype, which may be less perturbed by ETA-selective antagonists, with potential implications for PH treatment. Additionally, in multiple PH models, *Dhcr24^high^*tissue-remodeling macrophages demonstrated upregulated expression of *Csf1r*, a drug target of imatinib ^57^ and seralutinib ^58–60^, suggesting that these experimental therapies may act by reducing the pro-inflammatory effects of *Dhcr24^high^* macrophages. GSEA of PH target genes highlights the potential clinical relevance of these newly defined cell subtypes, based on their expression of target genes for existing and emerging PH therapies.

In conclusion, we have established single-cell profiles of multiple rodent models and integrated them with the HLCA, providing the first cross-species comparison of cell types and subtypes in experimental PH. We have identified new cell subtypes explaining the shared pathology and model-specific features. These novel populations were validated in patient tissues, confirming their relevance to human PH. Our comprehensive characterization of these rodent models and identification of the cells targeted by different therapeutic agents may facilitate future translational studies.

## Data availability

The datasets presented in this study can be found in online repositories. The name of the repository and accession number can be found at: https://ngdc.cncb.ac.cn/gsa/PRJCA009046. The single cell datasets of HLCA and PAH rats were downloaded from https://ega-archive.org/studies/EGAS00001004344 and http://mergeomics.research.idre.ucla.edu/PVDSingleCell/. The RNA microarray data of PAH patients was available at Gene Expression Omnibus (GEO) database as GSE117261.

### Author Contributions

B.L., P.Y., P.B.Y, J.W. and C.W. designed the study and interpreted the data. B.L., P.Y., Y.X. and Y.Z. drafted the manuscript. B.L., P.Y. and P.B.Y revised the manuscript. B.L., Y.Z., X.X. and J.P. analyzed the single cell data. Y.X. performed experiments. T.S. provided the Hx and SuHx models. X.W. provided the rat model. X.S. and H.Z. performed the single cell experiments.

## Acknowledgements

This study was supported by the Noncommunicable Chronic DiseasesNational Science and Technology Major Project (2024ZD0528700); National Natural Science Foundation of China (82241004, 82270062, Excellent Youth Scholars Program); Beijing Natural Science Foundation (Z220019); Beijing Outstanding Young Scientist Program (BJZQ2024002); Beijing Municipal Natural Science Foundation (7242096); National High Level of Hospital Clinical Research Funding (2022-PUMCH-D-002); Chinese Academy of Medical Sciences Innovation Fund for Medical Sciences (2023-I2M-2-001, 2021-I2M-1-049); Non-Profit Central Research Institute Fund of the Chinese Academy of Medical Sciences (2021-RC310-016); Haihe Laboratory of Cell Ecosystem Innovation Fund (22HHXBSS00010); State Key Laboratory Special Fund (2060204); Fundamental Research Funds for the Central Universities (3332023187).This work was supported with funding from the National Institutes of Health (NIH) National Heart, Lung and Blood Institute (R01HL159443). PBY is a co-founder and consultant for Keros Therapeutics, which develops therapies for cardiovascular, hematologic, and musculoskeletal diseases targeting bone morphogenetic protein and TGF-β signaling pathways. PBY is a co-founder of Modal Therapeutics, which develops therapies for vascular and metabolic diseases. PBY is a consultant for OrphAI Therapeutics, which develops therapies for pulmonary and vascular disease. PBY receives funding from and is co-inventor of intellectual property jointly owned with Pfizer, Inc. The interests of PBY are reviewed and managed by Massachusetts General Hospital in accordance with their conflict-of-interest policies.

